# Atypical scene-selectivity in the retrosplenial complex in individuals with autism spectrum disorder

**DOI:** 10.1101/2024.12.16.628702

**Authors:** Andrew S. Persichetti, Taylor L. Li, W. Dale Stevens, Alex Martin, Adrian W. Gilmore

## Abstract

A small behavioral literature on individuals with autism spectrum disorder (ASD) has shown that they can be impaired when navigating using map-based strategies (i.e., memory-guided navigation), but not during visually guided navigation. Meanwhile, there is neuroimaging evidence in typically developing (TD) individuals demonstrating that the retrosplenial complex (RSC) is part of a memory-guided navigation system, while the occipital place area (OPA) is part of a visually-guided navigation system. A key identifying feature of the RSC and OPA is that they respond significantly more to pictures of places compared to faces or objects – i.e., they demonstrate scene-selectivity. Therefore, we predicted that scene-selectivity would be weaker in the RSC of individuals with ASD compared to a TD control group, while the OPA would not show such a difference between the groups. We used functional MRI to scan groups of ASD individuals and matched TD individuals while they viewed pictures of places and faces and performed a one-back task. As predicted, scene-selectivity was significantly lower in the RSC, but not OPA, in the ASD group compared to the TD group. These results suggest that impaired memory-guided navigation in individuals with ASD may, in part, be due to atypical functioning in the RSC.

**Lay summary:** The retrosplenial complex (RSC), a cortical region that is part of a neural system that supports our ability to form map-like mental representations of the environment and use them to navigate (i.e., memory-guided navigation), exhibits atypical responses to images of places in individuals with autism spectrum disorder (ASD). These results are a first step towards understanding the neural mechanisms responsible for understudied behavioral impairments in memory-guided navigation in individuals with ASD.

## Introduction

The core impairments that define autism spectrum disorder (ASD) are related to social communication and restricted and repetitive behaviors (Marco et al., 2011; APA DSM-V, 2013). However, researchers have highlighted other cognitive domains that may be impaired in individuals with ASD, such as autobiographical memory retrieval and a lesser-known deficit in map-based (also referred to as “memory-guided” or “allocentric”) navigation – i.e., the ability to form map-like mental representations in memory and use them to navigate to out-of-sight places in the broader environment (Lind et al., 2013, 2014; Ring et al., 2018; Yang et al., 2021; for a recent review, see Agron et al., 2024). To date, however, reports of atypical navigation behaviors in ASD have not been accompanied by any reports of differences in neural activity associated with such behavioral impairments.

Several studies using functional MRI (fMRI) have demonstrated that the retrosplenial complex (RSC) and occipital place area (OPA) both respond significantly more to images of places (also called “scenes”) than to images of faces and objects, thus earning them the distinction of being scene-selective regions of human cortex (Maguire, 2001; Dilks et al., 2013). Beyond being scene-selective, it has been demonstrated that the RSC and OPA are parts of dissociable cortical systems that are involved in memory-guided navigation and visually guided navigation, respectively (Dilks et al., 2022). Since the RSC is part of a neural system that is responsible for the type of memory-guided navigation that is impaired in individuals with ASD, we used fMRI to ask whether processing of places or scenes might be atypical in a group of ASD participants. Specifically, we predicted that the RSC in individuals with ASD would show significantly weaker scene-selectivity when compared to a TD control group. In contrast, the OPA was not predicted to show between-groups differences, because evidence does not suggest differences in visually guided navigation behavior in individuals with ASD.

## Methods

### Participants

Twenty individuals [age, mean (SD) = 19.48 (2.9) years] who met the DSM-V criteria for ASD (APA DSM-V, 2013), as assessed by a trained clinician, were recruited for this experiment. All were high functioning and without intellectual disability. In addition, nineteen individuals with no history of psychiatric or neurological disorders [mean (SD) age = 21.01 (4.2) years] served as the TD control group (Some of the data from the TD group were previously described in Stevens et al., 2015,2017). There were no significant differences between the two groups in age (t_(37)_ = 1.31, p = 0.20) or overall IQ (Full-score IQ, mean (SD): ASD: 115.4 (12); TD: 119.3 (11.6), t_(37)_ = 1.04 p =0.31), which was measured using the Wechsler Abbreviated Scale of Intelligence (Wechsler, 1999) within one year of the scanning session in all participants. All participants from both groups were males with normal or corrected-to-normal visual acuity. Informed assent and consent were obtained from all participants and/or their parent/ guardian when appropriate, and all methods used in this study followed ethical guidelines and regulations in accordance with a National Institutes of Health (NIH) Institutional Review Board-approved protocol (10-M-0027, clinical trials number NCT01031407).

### MRI data acquisition and experimental task procedure

Scanning was completed on a General Electric Signa 3 Tesla scanner (GE Healthcare) with an 8-channel receive-only head coil at the NIH Clinical Center NMR Research Facility. For each participant, T2*-weighted blood oxygen level-dependent (BOLD) images covering the whole brain were acquired using a gradient echo single-shot echo planar imaging sequence (repetition time = 2,000 ms, echo time = 27 ms, flip angle = 77°, 41 axial contiguous interleaved slices per volume, 3.0-mm slice thickness, field of view = 216 mm, 72 × 64 acquisition matrix, single-voxel volume = 3.0 mm isotropic). In addition to the functional images, a high-resolution T1-weighted anatomical image (magnetization-prepared rapid acquisition with gradient echo—MPRAGE) was obtained (124 axial slices, 1.2mm^3^ single-voxel volume, 256 × 256 acquisition matrix, field of view = 24 cm).

During scanning, all participants completed six runs of a multicategory functional localizer with a block-design task. Each run lasted 7 m 18 s (219 brain volumes), and independent measures of nuisance physiological variables (cardiac and respiration) were recorded during all scans for later removal. Each run comprised fourteen task blocks (20 s each) interleaved with 13 fixation blocks (10 s each), with an additional fixation block at the beginning (18 s). There were fourteen different categories of stimuli used in the original study (for more details, see Stevens et al., 2015). However, we focused on places and faces (elliptically cropped to exclude hair and clothes) for the primary contrast in this report, with a supplemental analysis using places, tools, abstract objects, and non-manipulable objects. All stimuli were grayscale images of the same size (600 × 600 pixels) and were projected onto a screen behind the scanner and viewed via a mirror mounted to the head coil. Each block contained twenty pictures from the same category, with each picture presented for 300 ms, followed by a 500-ms interstimulus interval. To ensure attention to each stimulus, participants performed a one-back task, responding by pressing a button with their left index finger every time the same picture was presented twice in a row (this happened 1-2 times per block).

All data were preprocessed using the AFNI software package (Cox, 1996). First, the initial four TRs from each EPI scan were removed using 3dTcat to allow for T1 equilibration. Next, 3dDespike was used to bound outlying time points in each voxel within four standard deviations of the time series mean and 3dTshift was used to adjust for slice acquisition time within each volume (to t = 0). Each volume of the scan series was then aligned to the first retained volume of the scan using 3dvolreg. Finally, all scans were spatially blurred by a 6-mm Gaussian kernel (full width at half maximum) and divided by the mean of the voxelwise time series to yield units of percent signal change. To ensure that the fMRI data quality from both groups was matched, we computed the temporal signal-to-noise-ratio (tSNR) across the whole brain as well as a summary of in-scanner head motion using the @1dDiffMag program in AFNI. The groups did not differ in tSNR (t_(37)_ = 1.62, p = 0.11) or in-scanner head motion (t_(37)_ = 1.72, p = 0.10).

After preprocessing, we used data from two of the task runs to functionally define the RSC and OPA bilaterally as regions that responded significantly more to pictures of places than faces (i.e., that demonstrated scene-selectivity – Figure 1A). This was done separately in each participant. We then extracted the average response to each category during the remaining four runs from each region of interest (ROI) for further analysis. Thus, the data used to functionally define each ROI and the data used for subsequent analyses were independent of one another. Although the right RSC and OPA were defined in every participant, we could not define the left RSC or OPA in every participant: in the TD group, we defined 18/19 left RSC’s and 16/19 left OPA’s, while in the ASD group, we defined 15/20 left RSC’s and 19/20 left OPA’s. Therefore, we focused our analyses on the right hemisphere only.

**Figure 1.**
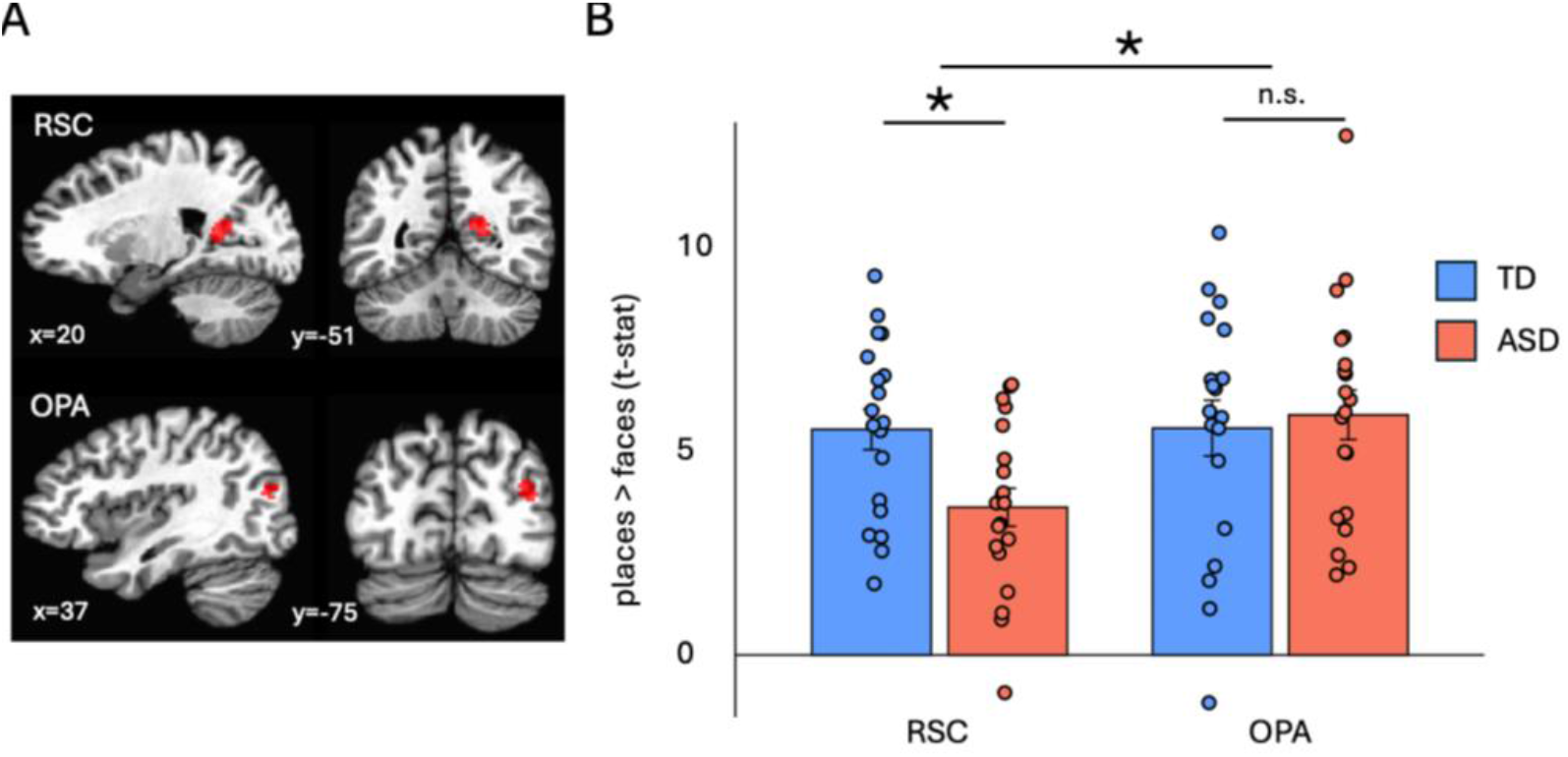
A) An example of the functionally defined RSC and OPA. The size and location of both regions of interest were very similar between the ASD and TD groups. B) A mixed-effects ANOVA showed that scene selectivity (places > faces) was weaker in the RSC, but not in the OPA, of the ASD group compared to the TD group.

## Results

First, the size and location of the RSC and OPA were compared between the TD and ASD participants. The sizes of each ROI did not significantly differ between the groups: the average number of 3mm^3^ voxels (in each participant’s native space) between the TD and ASD groups in the RSC was 9.58 and 11.05, respectively (t_(37)_ = 0.68, p = 0.50, Cohen’s d = 0.22), while the number of voxels in the OPA was 14.82 and 14.5, respectively (t_(37)_ = -0.08, p = 0.94, Cohen’s d = 0.03). Additionally, the Euclidean distances between the centers of mass for the average RSC and OPA across groups were less than one voxel’s width away from one another – 1.75 mm and 2.22 mm, respectively. Next, to test our hypothesis that scene-selectivity in ASD individuals is atypical in the RSC, but not the OPA, we ran a 2 Group (TD, ASD) x 2 ROI (RSC, OPA) mixed-effects ANOVA. As predicted, we found a significant Group x ROI interaction (F_(1,37)_ = 5.08, p = 0.03, ηp^2^ = 0.12) and independent-samples t-tests confirmed that scene-selectivity in the RSC of the ASD group was weaker than the TD group (t_(37)_ = 2.81, p < 0.01, Cohen’s d = 0.90), whereas there was no difference in scene-selectivity in the OPA between the groups (t_(37)_ = 0.36, p = 0.72, Cohen’s d = 0.12 – Figure 1B). A reasonable concern using this approach is that the use of faces as a comparison condition to scenes may be driving the apparent between-group difference, given that social deficits are a core impairment associated with ASD. We therefore repeated the above analysis using the average BOLD responses to viewing images of non-manipulable objects, tools, and abstract objects and used these responses, in turn, as contrasts to the responses to viewing images of places. In all cases, scene selectivity was weaker in the RSC of the ASD group (all t’s > 2.25, all p’s < 0.05).

## Discussion

In this report, we tested the hypothesis that impairments in memory-guided navigation in individuals with ASD may be due to atypical functioning in the RSC – a key node of the neural system underlying memory-guided navigation in humans. Our hypothesis was motivated by recent work suggesting that resting-state functional connectivity within a brain network that supports memory-guided navigation (and autobiographical memory retrieval), and includes the RSC, differs between groups of ASD and TD individuals (Persichetti et al., 2024). Furthermore, there is a fairly large literature of behavioral data demonstrating that mnemonic processes related to scene construction and autobiographical memory retrieval may be impaired in ASD (reviewed in Agron et al., 2024). Consistent with our predictions, we found that scene-selectivity was significantly lower in the RSC, but not in the OPA (which is instead involved in the distinct process of visually guided navigation) in a group of individuals with ASD compared to a tightly matched TD control group. These results are a first step towards understanding the neural mechanisms responsible for understudied impairments in memory-guided navigation in individuals with ASD.

In addition, the RSC has also been implicated in the ability to mentally generate and maintain a coherent representation of spatial contexts, or “scenes”, in the service of retrieving autobiographical memories (Hassabis et al., 2007). Thus, our finding of atypical RSC functioning in individuals with ASD might also shed light on well-documented difficulties in their ability to retrieve specific details associated with the recollection of autobiographical events (Kanner, 1943; Cooper and Simon, 2019; Agron et al., 2024). Therefore, the work described in this report is also a promising early step in understanding the neural systems that support memory-guided navigation and autobiographical memory retrieval, and how these distinct systems might overlap. Taken together, the results reported here address relatively understudied impairments related to ASD and thus may lead to a better understanding of the heterogeneous phenotypes observed in individuals with ASD.

## Acknowledgments

We thank Steve Gotts for insightful discussions and technical assistance. This work was supported by the NIMH Intramural Research Program. (#ZIA MH002920-09, clinical trials number NCT01031407). The authors declare no competing financial interests.

